# Aicardi-Goutières Syndrome Associated *ADAR* G1007R mutation dominantly induces neuroinflammation in mouse brain

**DOI:** 10.1101/2025.04.21.649787

**Authors:** Xinfeng Guo, Jia-Jun Liu, Chaowei Shang, Christopher J. Guerriero, Clayton Wiley, Richard A. Steinman, Yi Sheng, Jeffrey L. Brodsky, Timothy R. Billiar, Silvia Liu, Qingde Wang

**Author notes:** **Correspondence to:** Q Wang, University of Pittsburgh, BSTW943, 200 Lothrop Street, Pittsburgh, PA 15213; S Liu, University of Pittsburgh, Pittsburgh, PA 15213.

## Abstract

The *ADAR^G^*^*1007*^*^R^*mutation is one of the most frequent mutations found in type six Aicardi-Goutières Syndrome (AGS), a severe inflammatory encephalopathy in pediatric patients. We report here a mouse model bearing an human equivalent *ADAR^G^*^*1007*^*^R^* mutation, and the heterozygous mice recapitulated some pathologic features of *ADAR^G^*^*1007*^*^R^* AGS patients, including early-onset brain inflammation in heterozygous individuals and interferon-stimulated gene (ISG) expression within deep brain areas. Furthermore, we demonstrated that brain inflammation could be reversed by deletion of the cellular RNA receptor MDA5, which blocks the cellular RNA sensing signaling pathway. This model provides a unique tool for studying the molecular mechanisms underlying the heterozygous *ADAR^WT/G^*^*1007*^*^R^* mutation in AGS brain pathogenesis. It may also be a valuable platform for developing personalized therapies for patients with this specific mutation.

## Introduction

Adenosine deaminase acting on RNA 1 (ADAR1), encoded by *ADAR* in humans and *Adar i*n mice, is an RNA editing enzyme that carries out A-to-I RNA editing in double-stranded regions of cellular RNAs^1–3^. Mutations in the *ADAR* locus, together with mutations in eight other protein-coding genes, cause Aicardi-Goutières syndrome (AGS)^4,5^, a progressive autoinflammatory encephalopathy which often starts from early infancy or in children, and results in severe mental and physical disabilities^6–8^. The typical pathologic features include leukodystrophy, basal ganglia calcification, and excessive interferon (IFN) pathway activation in the brain, while multiorgan lesions similar to those in systemic lupus erythematosus (SLE) also occur in some AGS patients^5,7,9–11^. The mortality rate is generally high because there is no effective treatment to alter the course of this disease.

Dozens of mutations have been found in the *ADAR* locus in AGS patients^12–14^. These mutations, except for the single nucleoside guanosine to adenosine (G>A) alteration that results in the G1007R mutation, exhibit a recessive inheritance pattern and are present at homozygosity or in a biallelic genetic status in affected individuals. However, the G1007R mutation, one of the most frequent mutations in *ADAR*, causes neuronal pathogenesis when heterozygous^12–14^. The G1007R mutation resides in ADAR1’s catalytic RNA editing domain that mediates A-to-I editing, the most prevalent form of RNA editing in neuronal cells and a required function in embryogenesis.^15,16–20^—Neuronal pathogenesis associated with this mutation is therefore thought to be linked to deficient RNA editing activity^12^. Several other ADAR1 mutations, such as the K999N, Y1112F, and D1113H mutations, are also located in the catalytic domain, impacting ADAR1 RNA editing activity. However, these catalytic domain mutations are not sufficient for AGS pathogenesis when only one allele is affected in either human or mouse models^12–14,21,22^. It is unclear how an editing defect would lead to pathogenesis in individuals heterozygous in the G1007 mutation. Therefore, the unique effect of G1007R as a direct cause of pathogenesis is unexplained.

Due to limitations in accessing human brain tissues, especially at early disease stages as well as the scarcity of AGS patients, characterizing pathologic changes in the brain and determining the mechanisms underlying brain injury in AGS patients are challenging. Thus, animal models with AGS-causing mutations are necessary to understand the mechanisms underlying AGS neuroinflammatory pathogenesis and for therapeutic tests. However, most AGS mutations failed to recapitulate brain pathology and did not exhibit typical neurological phenotypes seen in AGS patients^23–27^. Nevertheless, deletion or mutations of the *Adar* gene in mice did cause activation of the IFN pathway, and some of those mutations result in inflammatory responses in the brain^22,28–30^. However, it is difficult to introduce a specific AGS-causing mutation at the ADAR1 catalytic domain due to high embryonic lethality^15,22,29^. A heterozygous mutation in the *Adar* locus or other AGS genetic loci has thus far not been demonstrated to trigger neuroinflammation.

We now report on the successful generation of a mouse model carrying an *Adar G956R* which is equivalent to the human *ADAR G1007R*; hereafter, we refer to this mutation as G1007R mutation. A G>A single nucleotide replacement was introduced into the mouse genome via CRISPR/Cas9. This mouse model replicated both the genetic and the neuroinflammatory responses seen in patients. Furthermore, we demonstrated that an MDA5-dependent RNA sensing signaling pathway plays an essential role in AGS neuroinflammation associated with the G1007R mutation, indicating that MDA5 might be a therapeutic target to treat *ADAR G1007R* AGS.

## Results

### 1. Heterozygous *Adar G1007R* mutation results in IFN pathway activation in the brain

The ADAR G1007R mutation, along with the K999N, Y1112F, and D1113H mutations found in AGS patients^12,13^, reside in the ADAR1 catalytic domain (**Figure 1A)**. However, only the G1007R mutation causes brain injury when present at heterozygousity^12–14^, suggesting a different mechanism of action associated with this specific mutation.

**Figure 1.**
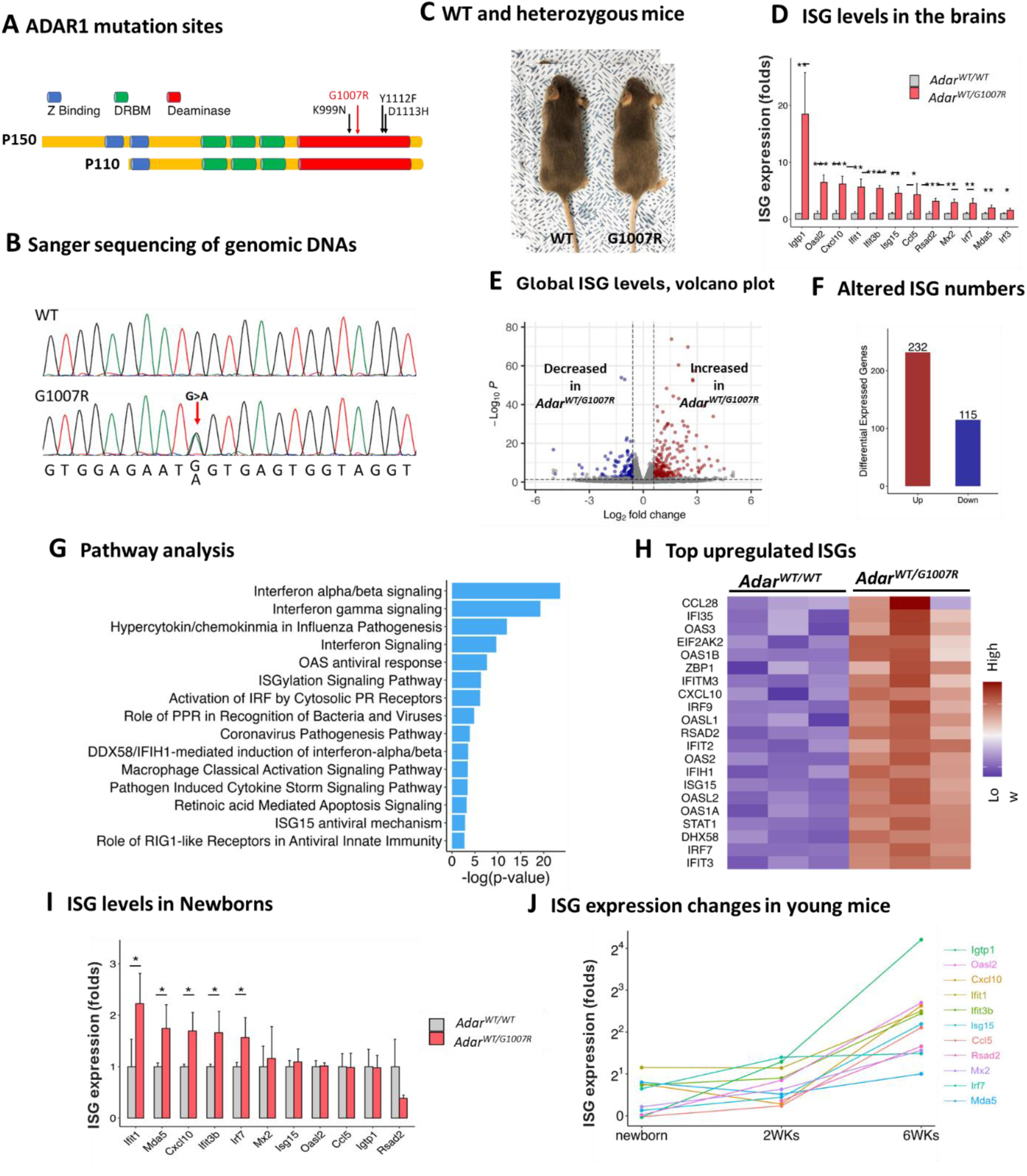
Phenotype of the heterozygous G1007R mutant mice. **A.** The *ADAR* gene encodes two isoforms of ADAR1 protein, p150 an p110. The Z binding domain, double-stranded RNA binding motifs (DRBM), and the deamination catalytic domains are shown in colors. The relative position of the G1007R mutation, together with other catalytic domain mutations found in AGS patients, are indicated by arrows. These catalytic mutations affect both the p150 and p110 isoforms of ADAR1. **B.** The genetic G>A single nucleotide mutation coding the G1007R amino acid replacement is confirmed by Sanger sequencing in the founder and progenies. This panel shows the chromatographs of the WT and heterozygous genomic DNA sequences flanking the mutation site. The G>A mutation site is indicated by the arrow. **C.** Heterozygous G1007R mutant mice do not exhibit abnormal phenotypes. The general development and body size are not different from the WT littermate control. Shown in this panel is a representative heterozygous mouse with comparison to its littermate at eight weeks of age. **D.** The brain ISG expression was assessed by real-time PCR, and the expression levels in *Adar^WT/G^*^*1007*^*^R^* heterozygous mice were compared to WT mice at the age of 6 weeks. All twelve tested ISG mRNAs were significantly increased in *Adar^WT/G^*^*1007*^*^R^* mice. Data are presented as mean ± SD. n=4 (WT), and n=5 (*Adar^WT/G^*^*1007*^*^R^*). The Prism t-test was used to test the differences between the two groups. * P<0.05, ** P<0.01,*** P<0.001. **E.** Global RNA expression level in the brains was assessed by RNA-seq analysis. The average values of three WT and three *Adar^WT/G^*^*1007*^*^R^* mice were shown in volcano plot. The significantly increased (red) and decreased (blue) genes in *Adar^WT/G^*^*1007*^*^R^* mice were highlighted. **F.** There are 347 genes in total transcript significantly changed expression levels in *Adar^WT/G^*^*1007*^*^R^* mice. The numbers of genes significantly upregulated or downregulated are shown. **G.** The genes with significantly changed expression were subjected to function analysis of Ingenuity Pathway Analysis (IPA). These genes fall into many functional pathways, especially those related to IFN signaling. The top 15 signaling pathway are show here. **H.** Interferon stimulated genes (ISG) are among the most significantly changed genes. The top 20 upregulated genes are show in this heat-map. **I.** ISG expression in the newborn brains is measured by real-time PCR. ISG expressions in the *Adar^WT/G^*^*1007*^*^R^* mice start to increase at newborn stage. Data are presented as mean ± SD. n=3 (WT), and n=5 (*Adar^WT/G^*^*1007*^*^R^*). The Prism t-test was used to test the differences between the two groups. * P<0.05. **J.** ISG expression levels in the brains of young *Adar^WT/G^*^*1007*^*^R^* mice were assessed at two different time points in addition to the newborns. ISG expressions were consistently increased from two weeks to six weeks. n=3 (WT), and n=5 (*newborn*), 4 (two weeks) and 5 (six weeks). All the tested ISGs are significantly increased in four and six weeks of age compared to the WT controls.

To divulge the molecular mechanism underlying the *ADAR G1007R* mutation in neuroinflammatory pathogenesis, we decided to generate a mouse model carrying the *G1007R* mutation. To this end, we used CRISPR/Cas9 to introduce a single G>A nucleotide replacement, equivalent to the *G1007R* mutation found in AGS patients, into the mouse *Adar* locus at the corresponding site, and eventually identified a founder carrying the designed mutation. We confirmed via Sanger sequencing that the founder and its progeny carried the G>A mutation (**Figure 1B)**.

We then asked whether the heterozygous *Adar* mutation (*Adar^WT/G^*^*1007*^*^R^*) causes phenotypes seen in humans. The disease onset of *ADAR^WT/G^*^*1007*^*^R^*AGS patients varies significantly, ranging from presentation at a neonatal stage, frequently associated with progressive and severe neural degeneration^12,13,31^, to presentation in young adults, often showing mild and relatively stable clinical signs^6,7,11,32,33^. The heterozygous *Adar^WT/G^*^*1007*^*^R^* mice did not exhibit an apparent neurologic phenotype that includes an exaggerated startle response, abnormal gait, muscle spasm, dystonia, or seizures (**Figure 1C)**. Decreased survival and growth retardation were also absent. Thus, this *ADAR^WT/G^*^*1007*^*^R^* model does not resemble the severe, early onset phenotype of AGS.

Next, we examined whether the G1007R mutation causes a latent stage or a mild AGS phenotype as seen in late-onset AGS patients. We assessed the inflammation levels in the brain by measuring the levels of a panel of twelve selected interferon stimulated gene (ISG) mRNAs in whole brain RNA samples isolated from six-week-old (young adult stage) *Adar^WT/G^*^*1007*^*^R^* heterozygous mice by real-time PCR. All tested ISGs exhibited dramatically increased expression levels compared to the littermate WT controls (**Figure 1D)**.

For a complete assessment of altered gene expression, we carried out a whole transcriptome RNA sequencing study (RNA-seq). Total brain RNAs from three WT mice and three heterozygous *Adar ^WT/G^*^*1007*^*^R^* mice were isolated and subjected to RNA-seq analysis. Among the whole transcriptome, 232 genes were expressed at significantly higher levels in *Adar ^WT/G^*^*1007*^*^R^*mice by comparison to the WT controls, while 115 genes showed decreased expression (**Figure 1E, F).** IPA pathway analysis showed that these differentially expressed genes were predominantly associated with innate immune functions, especially the IFN signaling pathways (**Figure 1G).** The top 20 significantly activated IFN pathway genes are shown in **Figure 1H**.

### 2. ISG expression rapidly increases in young *Adar ^WT/G^*^*1007*^*^R^* mice and is predominantly expressed in deep brain areas

Limited by the accessibility of brain tissues from AGS patients, it is difficult to determine the innate immune response in the brain, especially at early disease stages. In a late-onset AGS patient, it is unknown if inflammation is initiated at the newborn stage or whether it starts at the time of disease onset at an older age. The *Adar^WT/G^*^*1007*^*^R^* mutant mice enabled us to determine the time course of inflammation development in the brain. Therefore, we measured ISG levels at birth, two weeks, four weeks, and six weeks. We found that the levels of *Ifit1, Mda5, Ifit3b,* and *Irf7* were significantly higher in newborn mice (**Figure 1I)**. We also found that ISG levels remain at a relative low level until two weeks of age, then consistently increased at four and six weeks of age (**Figure 1J).** This finding indicates that low level neuroinflammation may present in patients preceding the onset of disease in the asymptomatic stage.

Next, we investigated whether the brain image features observed in AGS patients correlate with the pathological pattern of ISG expression in *Adar ^WT/G^*^*1007*^*^R^*mice. We carried out an RNA i*n situ* hybridization study to determine the location of ISG expression in brain sections of *six-weeks old* mice. We used four representative RNAScope^TM^ probes to detect *Ifit1, Isg15, Cxcl10 and Oasl2* expression. These probes detected substantial ISG expression in the *Adar ^WT/G^*^*1007*^*^R^* brains while no obvious ISG signal was detected in the WT controls (**Figure 2A, B)**. Nevertheless, the ISG signals were not evenly distributed throughout the brains but were predominantly present in ependymal and deep brain regions, especially in periventricular areas (**Figure 2A, B and Figure 3A)**. Three probes, *Cxcl10, Isg15 and Oasl2* showed a similar staining pattern, and extensive hybridization was seen in ependymocytes and in cells with neuronal morphology adjacent to the ventricles. The *Cxcl10* probe stained a more limited number of ependymocytes. The *Ifit1* probe showed stronger staining of the periventricular rather than ependymal cells, while the *Isg15, Cxcl10 and Oasl2* probes equally or preferentially stained the ependymocytes. In the cortex (CTX) areas, scattered cell clusters were positively stained by the four probes (**Figure 3A)**. Serial sections demonstrated that scattered cell clusters consistently showed staining for all four probes. The *Ifit1* and *Oasl2* probes stained more cells, while the *Cxcl10* stained the fewest. These ISG staining patterns suggested that different regulatory mechanisms underlie ISG expressions in different brain cells. Next, we investigated whether increased ISG expression leads to neuroinflammatory responses detectable through morphological analysis. We assessed astrocytosis and microgliosis in the brain using immunohistochemistry studies with antibodies specific for glial fibrillary acidic protein (GFAP) and ionized calcium binding adaptor molecule 1 (Iba1), markers for astrocytosis and microgliosis, respectively. Brain tissue from three WT and three *Adar ^WT/G^*^*1007*^*^R^* mice at six weeks of age was serially sectioned and stained with these antibodies. We observed that GFAP and Iba1 staining was more intense in the periventricular areas of *Adar ^WT/G^*^*1007*^*^R^* mice than that in WT mice. compared to WT mice (**Figure 3B**). Quantitative analysis of 20–30 measured regions in the WT and *Adar ^WT/G^*^*1007*^*^R^*brain sections showed that the average signal intensity of both markers was significantly higher in *Adar ^WT/G^*^*1007*^*^R^* mice than in WT mice (**Figure 3C**). This suggests that the mutation-triggered innate immune activation leads to neuroinflammatory responses in the brains of heterozygous G1007R mice. Therefore, the ISG expression pattern in deep brain regions, along with increased astrocytosis and microgliosis in *Adar ^WT/G^*^*1007*^*^R^* mice, well correlates with the pathologic features observed in brain images of AGS patients^12,13^.

**Figure 2.**
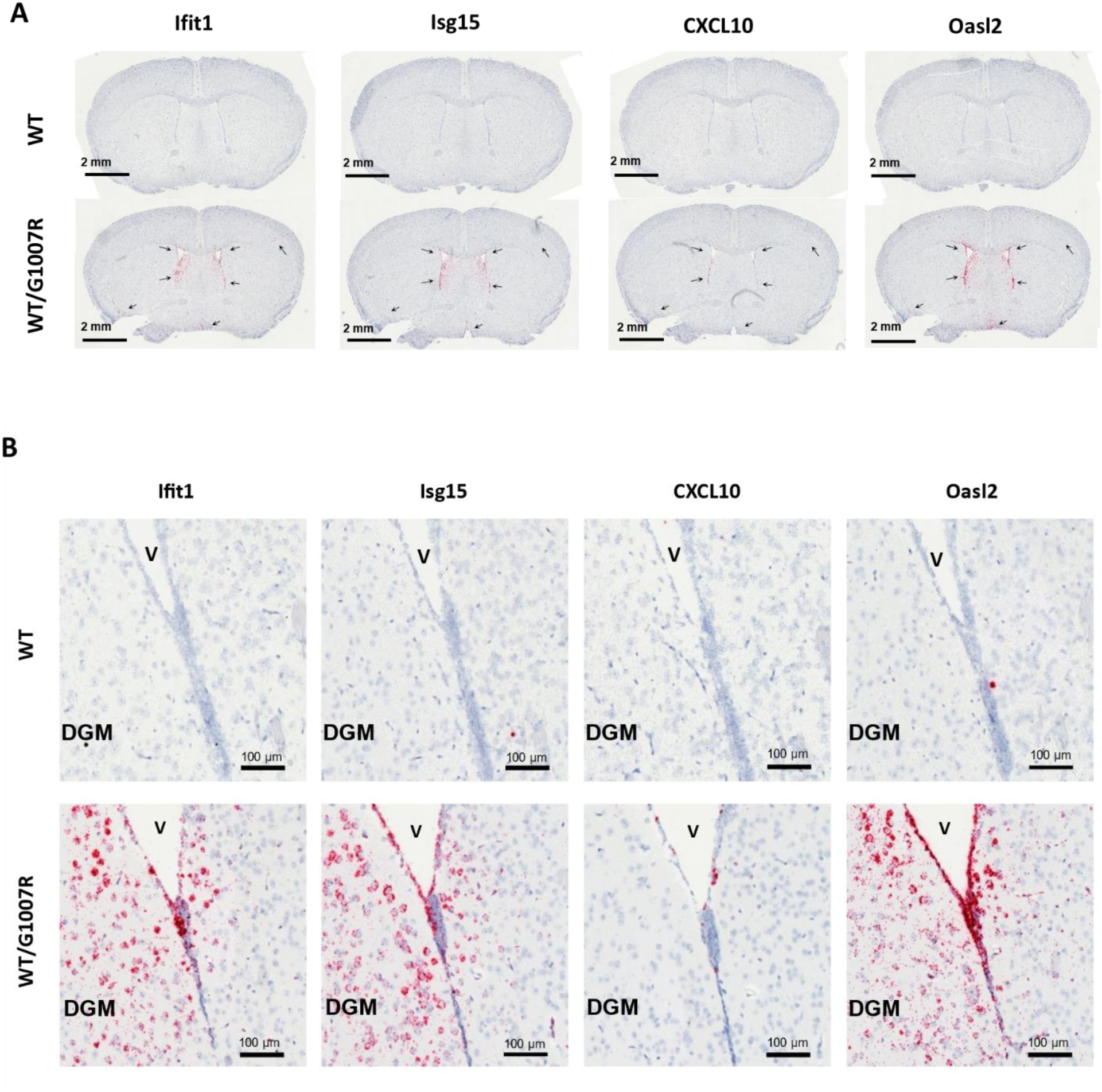
ISG expression pattern in the brain of heterozygous *Adar^WT/G^*^*1007*^*^R^* mice. **A.** ISG expression on coronally sectioned WT and *Adar^WT/G^*^*1007*^*^R^* mouse brains are detected by *in situ* hybridization (ISH) with RNAscope probes for *Ifit1, Isg15, Cxcl10 and Oasl2*. Digital images of brain staining are collected by whole slide image scan. The periventricular areas show strong ISG signals, indicated by the arrows. Scatted cell clusters in the cortex areas are also observed. No signal is observed in WT brains. Scale bar =2mm. **B.** The ISG expression (red) in deep gray matter and the periventricular regions is shown in high power images. While no signal is observed in WT brains, *Ifit1, Isg15 and Oasl2* probes show strong staining in cells with neuronal morphology in periventricular regions. Ependymocytes show intense hybridization for *Isg15, Ccxl10 and Oasl2* probes. The *Cxcl10* probe stains fewer cells, and the distribution and morphology of *Cxcl10* positive cells is consistent with ependymal cell type. Bar scale is 100um as indicated. V, ventricle.

**Figure 3.**
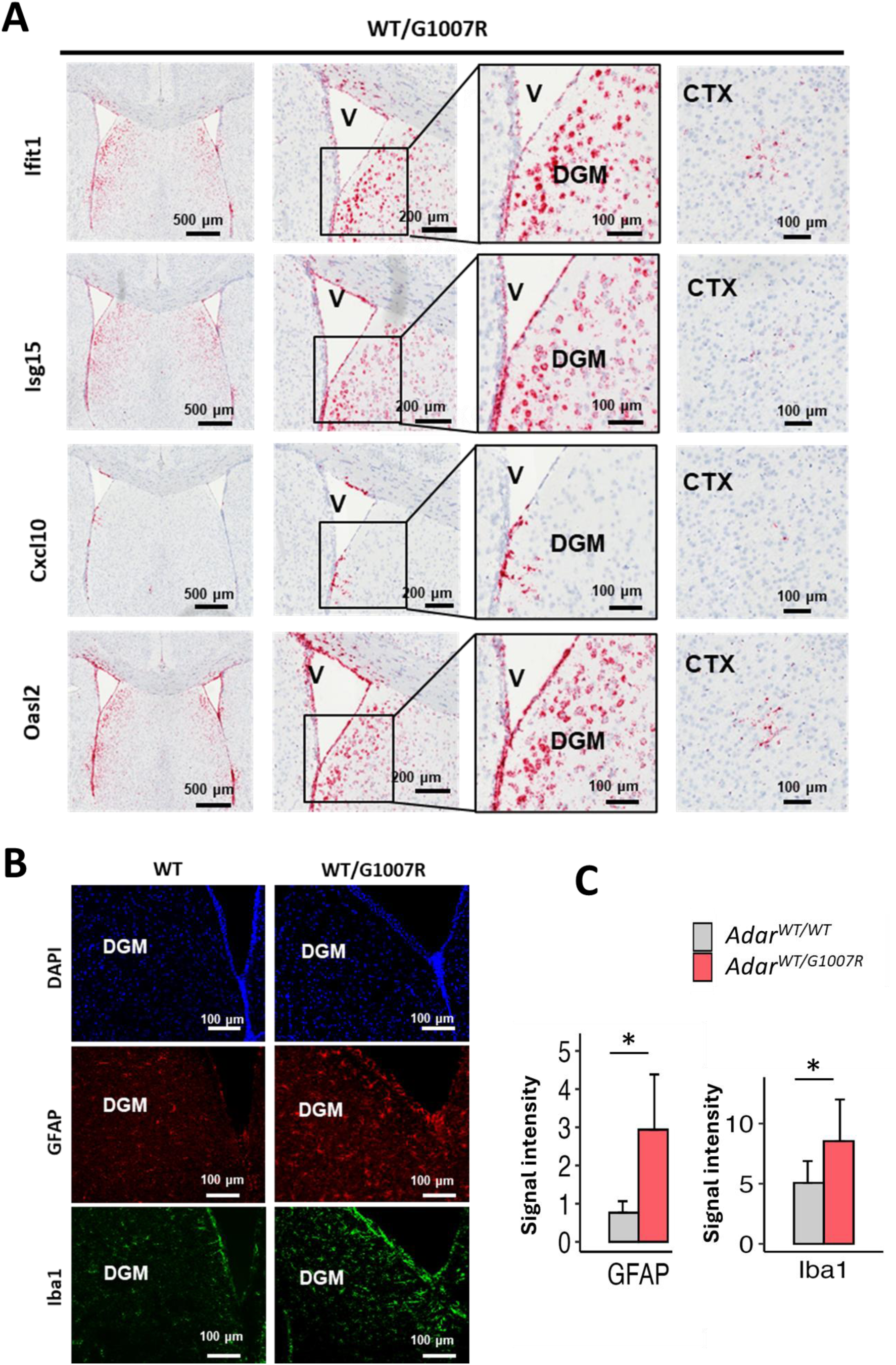
ISG expression and inflammatory response in deep brain areas of *Adar^WT/G^*^*1007*^*^R^* mice. **A.** The ISG expression patterns as shown by *in situ* hybridization (red) of the indicated probes in periventricular (first 3 columns) and cortical (4th column) regions of *Adar^WT/G^*^*1007*^*^R^* mice are shown in low (1st and 2nd column) and high (3rd and 4th column) power images. Bar scales are 500um, 200um and 100um as indicated. V, ventricle. DGM, deep gray matter, CTX, cortex. *Ifit1, Isg15 and Oasl2* probes show strong staining in cells with neuronal morphology in periventricular regions. Ependymocytes show intense hybridization for *Isg15, Ccxl10 and Oasl2* probes. The *Cxcl10* probe stains fewer cells in both periventricular and cortical regions. The distribution and morphology of *Cxcl10* positive cells is consistent with ependymal and microglial cell types. **B.** Immunofluorescence microscopy of cryosections from WT and *Adar^WT/G^*^*1007*^*^R^* mouse brains stained with GFAP (astrocytes) and Iba1 (microglia). Increased GFAP and Iba1 staining intensity is observed in the periventricular areas of *Adar^WT/G^*^*1007*^*^R^* brains compared to WT controls. DGM, deep gray matter. Scale bar =100um. **C.** Quantification of GFAP and Iba1 fluorescence intensities in WT and *Adar^WT/G^*^*1007*^*^R^* mouse brains. Both markers are significantly elevated in *Adar^WT/G^*^*1007*^*^R^* brains. Data are presented as mean ± SD, with measurements collected from nine sections from three brains per group, including 20 (WT) and 30 (*Adar^WT/G^*^*1007*^*^R^*) measured areas. Statistical significance was determined using an unpaired *t*-test via Prism software. * P<0.05.

Together, the heterozygous *Adar^WT/G^*^*1007*^*^R^* mutation is sufficient to induce significant ISG expression and inflammatory response in mouse brains, recapitulating genetic and pathologic features of AGS patients. However, the *Adar^WT/G^*^*1007*^*^R^* mouse model did not recapitulate the full degenerative phenotype described in severe early-onset AGS patients but more so a latent stage of late-onset AGS.

### 3. *Adar^WT/G^*^*1007*^*^R^* mutation causes innate immune activation through MDA5-dependent RNA sensing signaling

MDA5 is a cytosolic dsRNA receptor that detects viral or endogenous RNAs to activate the innate immune system, leading to type I interferon expression^34–36^. When the integrity of either the RNA editing activity or the Z-RNA binding activity of ADAR1 is impaired, cellular RNAs stimulate MDA5 to activate the innate immune response^15,22,29,30,37–40^. To test the hypothesis that the ADAR1 G1007R mutation causes neuroinflammation in the brain of *Adar ^WT/G^*^*1007*^*^R^* mice through the MDA5-dependent RNA sensing signaling pathway, we crossed *Adar^WT/G^*^*1007*^*^R^* and *Ifih1^−/-^*(MDA5 KO) mice^41^ to produce *Adar^WT/G^*^*1007*^*^R^*; *Ifih1^−/-^* double mutant mice. We then analyzed ISG expression levels in the brains of *Adar^WT/G^*^*1007*^*^R^*; *Ifih1^−/-^* mice by real-time PCR. All eight tested genes were dramatically decreased in the *Adar^WT/G^*^*1007*^*^R^*; *Ifih1^−/-^* mice compared to *Adar^WT/G^*^*1007*^*^R^*mice, and in fact the levels were now comparable to the WT control mice; the expression of *Mx2* was even lower than the WT controls (**Figure 4A)**. We also performed an RNA-seq study on *Adar^WT/G^*^*1007*^*^R^*; *Ifih1^−/-^* mice and compared the gene expression signature with WT and *Adar^WT/G^*^*1007*^*^R^* mice. Among the 232 upregulated genes observed in *Adar^WT/G^*^*1007*^*^R^*mice, 126 genes were in the IFN signaling pathways, and 109 of these genes were dramatically decreased in the *Adar^WT/G^*^*1007*^*^R^*; *Ifih1^−/-^* mice to the level of control mice. Only 17 of the 126 genes remained at the levels of the *Adar^WT/G^*^*1007*^*^R^*mice (**Figure 4B).** These results indicate that IFN pathway activation in the *Adar^WT/G^*^*1007*^*^R^* brains was largely dependent on the MDA5-RNA sensing signaling pathway. As expected, MDA5 deletion did not change the expression of the remaining 106 up- and 115 down-regulated non-IFN stimulated genes in *Adar^WT/G^*^*1007*^*^R^*; *Ifih1^−/-^* mice (**Figure 4C)**. Pathway analysis also showed that these non-IFN stimulated genes were associated with multiple functions, including extracellular matrix organization, cell junction organization, degradation of the extracellular matrix, and others (**Figure 4D).** Since the *Adar^WT/G^*^*1007*^*^R^*; *Ifih1^−/-^* mice did not exhibit any obvious phenotypes, the role of these genes is likely insignificant.

**Figure 4.**
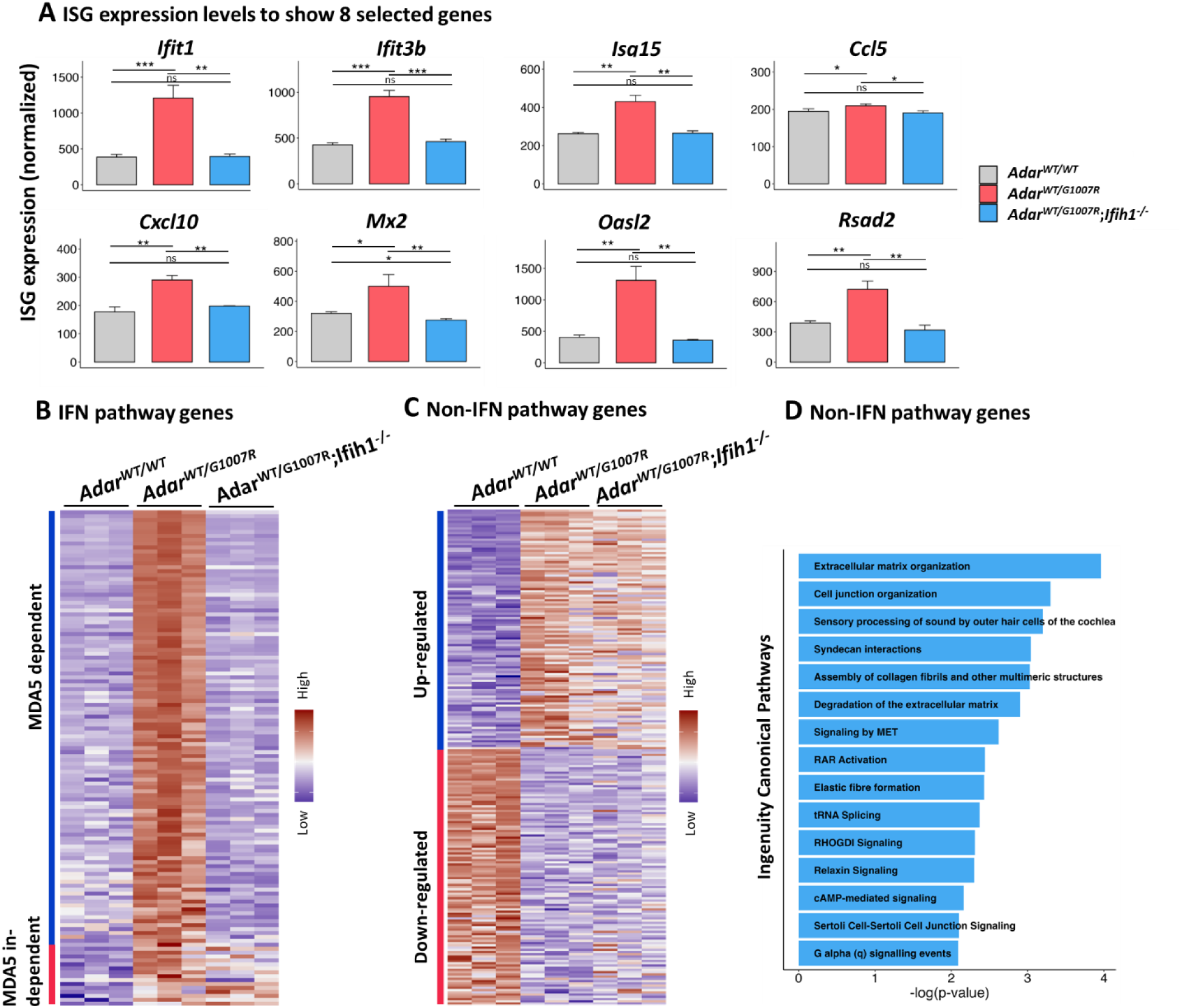
MDA5-dependent and independent ISG expressions caused by G1007R mutation. A. ISG expression in the brains of *Adar^WT/G^*^*1007*^*^R^*; *Ifih1^−/-^* mice are assessed by real-time PCR and compared with WT and *Adar^WT/G^*^*1007*^*^R^*mice. As shown by the eight selected ISGs, MDA5 deletion in the *Adar^WT/G^*^*1007*^*^R^*; *Ifih1^−/-^*mice dramatically suppressed ISG expression in the brain. The ISG levels in *Adar^WT/G^*^*1007*^*^R^*; *Ifih1^−/-^*mice are significantly decreased compared to *Adar^WT/G^*^*1007*^*^R^* mice, and there is no significant difference from the WT mice. Data are presented as mean ± SD, n=4 (WT), and n=5 (*Adar^WT/G^*^*1007*^*^R^* and *Adar^WT/G^*^*1007*^*^R^*; *Ifih1^−/-^*). The 1-way ANOVA test was used to test the differences between the WT and *Adar^WT/G^*^*1007*^*^R^* , *Adar^WT/G^*^*1007*^*^R^* and *Adar^WT/G^*^*1007*^*^R^*; *Ifih1^−/-^* , and WT and *Adar^WT/G^*^*1007*^*^R^* groups. * P<0.05, ** P<0.01,*** P<0.001. B. Heatmap of ISG expression levels in the WT, *Adar^WT/G^*^*1007*^*^R^* and *Adar^WT/G^*^*1007*^*^R^*; *Ifih1^−/-^* mice. Among the differentially expressed genes (DEGs) shown by RNA-seq analysis (Figure 1E), expression of 126 ISGs are significantly upregulated in *Adar^WT/G^*^*1007*^*^R^* mice, while 109 upregulated genes are reduced to WT levels in the *Adar^WT/G^*^*1007*^*^R^*; *Ifih1^−/-^* mice (MDA5 dependent). While 17 upregulated ISGs in *Adar^WT/G^*^*1007*^*^R^* mice do not decrease their expression in *Adar^WT/G^*^*1007*^*^R^*; *Ifih1^−/-^* mice (MDA5 independent). C. Heatmap of the non-IFN pathway genes in the WT, *Adar^WT/G^*^*1007*^*^R^* and *Adar^WT/G^*^*1007*^*^R^*; *Ifih1^−/-^* mice. Bothe up and down regulated genes in *Adar^WT/G^*^*1007*^*^R^* mice do not change expression with deletion of MDA5 in the *Adar^WT/G^*^*1007*^*^R^*; *Ifih1^−/-^* mice. D. Functional analysis of the non-IFN pathway genes which expressions are significantly changed in *Adar^WT/G^*^*1007*^*^R^* mice, either upregulated or downregulated. IPA pathway analysis shows that these genes are associated with extracellular matrix organization, cell junction organization, degradation of the extracellular matrix, and others.

## Discussion

The *ADAR* G1007R mutation is one of the most frequent mutations in type 6 AGS, an inflammatory encephalopathy caused by mutations in *ADAR* locus^12–14^. However, the molecular mechanism underlying this disease is poorly understood. To date, *in vivo* studies on AGS have been restricted by a lack of appropriate animal models that recapitulate the genetic and inflammatory features associated with the disease, most notably in the brain. In this study, we successfully generated a mouse model that carries a single nucleotide mutation equivalent to the patient *ADAR* G1007R mutation. Like in patients, the mutation resulted in brain inflammation in mice when it presented on only one chromosome. We also found that the *Adar* G1007R mutation altered RNA splicing patterns in ADAR1-associated transcripts that result in protein depletion, thereby dramatically impacting the RNA editing function to trigger infoammation. We further demonstrated that the *Adar* G1007R mutation causes brain inflammation through the MDA5-dependent RNA sensing signaling pathway.

In patients, early-onset AGS usually exhibit severe brain injury in the first few weeks of life, while late-onset AGS is associated with normal development for months, years, or even decades until symptoms appear that often follow viral infection or vaccination^12,13^. While the *Adar^WT/G^*^*1007*^*^R^* mice did not exhibit an obvious neurological phenotype, an innate immune was activated in the brain. Despite the increase in ISG expression in the brain, the typical neurological phenotypes observed in patients did not present in other AGS mouse models either^22,28–30^. Differences in genomic sequence between humans and mice may have contributed to the different phenotypes, e.g. there are significantly fewer A-to-I RNA editing substrates in mice than in humans^42–44^. Thus, an ADAR1 mutation is likely more deleterious in human cells because more unedited RNAs are produced, which would trigger a severer immune response. Nevertheless, the *Adar* G1007R mutation resulted in robust ISG expression in the heterozygous mice, and the location of ISG expression in the brain was consistent with the pathologic features observed in the brains of AGS patients. Whether the pathogen-free environment in which the mice were housed contributed to the phenotype of *Adar* G1007R mice has yet to be determined but will be an important focus for future studies. It is currently impossible to determine whether elevated ISG expression in the human brain precedes the onset of AGS, but the results from mice of this study provide clues for the pathogenesis of the AGS brain caused by the *ADAR^WT/G^*^*1007*^*^R^*mutation. Finally, we demonstrated that the homozygous *Adar* G1007R mutation is embryonic lethal, explained why only the heterozygous *ADAR* G1007R mutation is found in AGS patients.

In sum, we demonstrate that heterozygous *Adar* G1007R mutation causes MDA5-dependent RNA sensing signaling pathway activation. MDA5 deletion prevented ISG expression in the brains of heterozygous mice. This finding indicate that the IFN pathway—and more specifically the cellular RNA sensor MDA5—is a potential therapeutic target for *Adar* G1007R-associated disease. This *Adar* G1007R mutant mouse model generated in this study therefore provides a critical tool to test AGS therapeutics and determine their mechanisms of action.

## Material and Method

### 1. Mouse genome mutagenesis

All mice used in this study were kept in a specific pathogen-free animal facility and all studies were carried out following protocols approved by our Institutional Animal Care and Use Committee. The G1007R mutation was introduced into the mouse genome using our CRISPR/Cas9 protocol as described previously^22,29,45,46^. Considering the genetic diversity of human diseases of ADAR1 mutations, mice used in this study, including *Adar^G^*^*1007*^*^R/G^*^*1007*^*^R^* and *Adar^G^*^*1007*^*^R/G^*^*1007*^*^R^*; *Ifih1^−/-^* mice, were purposely maintained in C57/BL6J and DBA2 mixed background. The sequence of the sgRNA: 5’-gcaaggcaagcttcgcacca-3’. The ssODN template sequence for the G to A mutation:5’-gtgccgtggaaagcacagagtcccgccattaccctgtctttgaaaatcccaagcaaggcaagcttcgcaccaaagtggagaatagggagtggtaggtgccagctggcagtgaggagacatgcacgcgaggggtgtccgcttcctt-3’. The primer sequences used for amplifying the region flanking G1007R mutation: 5’-tgccagttcccacataggat-3’ and 5’-agtccagtgacacccacctc-3’.

Founders carrying the single G>A nucleotide replacement were bred to homozygosity, and bred with *Ifih1^−/-^* mice for phenotypic analysis. *Ifih1^−/-^* mice were purchased from Jackson lab, strain #: 015812.

### 2. ISG expression quantification

RNA isolation was performed with RNeasy Plus Mini Kit (Qiagen Cat # 74134) following the manufacturer’s instructions. Quantitative RT-PCR was performed using the iTaqTM Universal SYBR Green One-Sep Kit (Bio-Rad cat #1725151). Assayed genes comprised *Cxcl10*, *Igtp1*, *Ifit1*, *Ccl5*, *Oasl2*, *Mx2*, *Isg15*, *Rsad2*, *Irf7*, and *Ifih1. Gapdh* and *Hprt* were used as internal controls. The specificity of PCR amplifications was confirmed by the melting curve and by electrophoresis analysis of the final PCR products. The quantification of the mRNA levels was calculated by the Ct values using ΔΔt method with internal references of the average value of the *Hprt* and *Gapdh* measurements as described previously^21,22^.

### 3. RNA *in situ hybridization* and Pathology studies

Mouse brains were fixed with 10% neutral buffered formalin and paraffin embedded tissues (FFPE) were sectioned at 6um thickness. H-E staining and RNA *in situ hybridization* (ISH) were performed on FFPE sections. Four commercial RNAscope Target Probes were purchased from Advanced Cell Diagnostics, Hayward, CA. The Ifit1 probe, cat # 500071, targets the sequence of 739-1846 of NM_008331.3; the ISG-15 probe, cat # 559271, targets the sequence of 2-561 of NM_015783.3; the CXCL10 probe, cat # 408921 targets the sequences of 11-1012 of NM_021274.2, and the Oasl2 probe, cat # 534501, target the sequence of 1201-2184 of NM_011854.2. FFPE section pretreatment, hybridization and signal detection were performed using the RNAscope™ 2.5 HD Detection Reagents-RED kit, cat # 322360 according to manufacturer’s protocols and as we previously described^22,29,30^. The stained brain sections were scanned with Fritz Precipoint digital scanner with 20X objective lens for image analysis.

### 4. Statistics analysis

Prism-GraphPad was used to conduct statistical data analysis and t-tests were used for ISG expression, RNA editing efficiency and body weight comparisons. RNA-seq data was analyzed and visualized using R programming with the available R/Bioconductor packages.

## Acknowledgements

We thank the University of Pittsburgh Biospecimen Core for brain tissue pathologic processing. We thank the University of Pittsburgh Center for Biologic Imaging for assistance in pathologic imaging.

This study was supported by the National Institutes of Health Grant R01AI139544, R01 NS134651 and R01 NS138573, as well as the University of Pittsburgh Brain Institute Assault on Alzheimer’s seed grant program. J.L.B. acknowledges support from National Institutes of Health Grant R35GM131732. This study was supported in part by the University of Pittsburgh Center for Research Computing sing the HTC cluster, which is supported by NIH award S10OD028483.

## Author contributions

XG carried out all animal model preparations and biochemical and molecular analysis; JL performed bioinformatics analysis on RNA-seq data and figure preparations. CG performed protein modeling studies. RS and JLB participated in experimental design, data analysis, and editing of the manuscript. YS participated in mouse model preparation; TB participated in experimental data discussion, data analysis, and manuscript writing; SL performed bioinformatics analysis on RNA-seq data and organized data presentation. QW designed the project, supervised the study, and wrote the manuscript.

All authors read and approved the final manuscript.

## References

1 Song, B., Shiromoto, Y., Minakuchi, M. & Nishikura, K. The role of RNA editing enzyme ADAR1 in human disease. Wiley Interdiscip Rev RNA 13, e1665, doi:10.1002/wrna.1665 (2022).

2 Wang, Q., Li, X., Qi, R. & Billiar, T. RNA Editing, ADAR1, and the Innate Immune Response. Genes (Basel) 8, doi:10.3390/genes8010041 (2017).

3 Lamers, M. M., van den Hoogen, B. G. & Haagmans, B. L. ADAR1: “Editor-in-Chief” of Cytoplasmic Innate Immunity. Front Immunol 10, 1763, doi:10.3389/fimmu.2019.01763 (2019).

4 Uggenti, C. et al. cGAS-mediated induction of type I interferon due to inborn errors of histone pre-mRNA processing. Nat Genet 52, 1364–1372, doi:10.1038/s41588-020-00737-3 (2020).

5 Crow, Y. J. et al. Characterization of human disease phenotypes associated with mutations in TREX1, RNASEH2A, RNASEH2B, RNASEH2C, SAMHD1, ADAR, and IFIH1. Am J Med Genet A 167A, 296–312, doi:10.1002/ajmg.a.36887 (2015).

6 Goutieres, F., Aicardi, J., Barth, P. G. & Lebon, P. Aicardi-Goutieres syndrome: an update and results of interferon-alpha studies. Ann Neurol 44, 900–907, doi:10.1002/ana.410440608 (1998).

7 Crow, Y. J. in GeneReviews((R)) (eds M. P. Adam et al.) (1993).

8 Aicardi, J. & Goutieres, F. A progressive familial encephalopathy in infancy with calcifications of the basal ganglia and chronic cerebrospinal fluid lymphocytosis. Ann Neurol 15, 49–54, doi:10.1002/ana.410150109 (1984).

9 Crow, Y. J., Vanderver, A., Orcesi, S., Kuijpers, T. W. & Rice, G. I. Therapies in Aicardi-Goutieres syndrome. Clin Exp Immunol 175, 1–8, doi:10.1111/cei.12115 (2014).

10 Crow, Y. J. Aicardi-Goutieres syndrome. Handb Clin Neurol 113, 1629–1635, doi:10.1016/B978-0-444-59565-2.00031-9 (2013).

11 Rice, G. et al. Clinical and molecular phenotype of Aicardi-Goutieres syndrome. Am J Hum Genet 81, 713–725, doi:10.1086/521373 (2007).

12 Rice, G. I. et al. Mutations in ADAR1 cause Aicardi-Goutieres syndrome associated with a type I interferon signature. Nature genetics 44, 1243–1248, doi:10.1038/ng.2414 (2012).

13 Rice, G. I. et al. Genetic, Phenotypic, and Interferon Biomarker Status in ADAR1-Related Neurological Disease. Neuropediatrics 48, 166–184, doi:10.1055/s-0037-1601449 (2017).

14 Vanderver, A. et al. Janus Kinase Inhibition in the Aicardi-Goutieres Syndrome. N Engl J Med 383, 986–989, doi:10.1056/NEJMc2001362 (2020).

15 Liddicoat, B. J. et al. RNA editing by ADAR1 prevents MDA5 sensing of endogenous dsRNA as nonself. Science 349, 1115–1120, doi:10.1126/science.aac7049 (2015).

16 Bazak, L. et al. A-to-I RNA editing occurs at over a hundred million genomic sites, located in a majority of human genes. Genome research 24, 365–376, doi:10.1101/gr.164749.113 (2014).

17 Levanon, E. Y. et al. Systematic identification of abundant A-to-I editing sites in the human transcriptome. Nature biotechnology 22, 1001–1005 (2004).

18 Kim, D. D. et al. Widespread RNA editing of embedded alu elements in the human transcriptome. Genome research 14, 1719–1725 (2004).

19 Blow, M., Futreal, P. A., Wooster, R. & Stratton, M. R. A survey of RNA editing in human brain. Genome research 14, 2379–2387 (2004).

20 Athanasiadis, A., Rich, A. & Maas, S. Widespread A-to-I RNA editing of Alu-containing mRNAs in the human transcriptome. PLoS biology 2, e391 (2004).

21 Guo, X. et al. ADAR1 RNA editing regulates endothelial cell functions via the MDA-5 RNA sensing signaling pathway. Life Sci Alliance 5, doi:10.26508/lsa.202101191 (2022).

22 Guo, X. et al. Aicardi-Goutieres syndrome-associated mutation at ADAR1 gene locus activates innate immune response in mouse brain. J Neuroinflammation 18, 169, doi:10.1186/s12974-021-02217-9 (2021).

23 Morita, M. et al. Gene-targeted mice lacking the Trex1 (DNase III) 3’-->5’ DNA exonuclease develop inflammatory myocarditis. Mol Cell Biol 24, 6719–6727, doi:10.1128/MCB.24.15.6719-6727.2004 (2004).

24 Mackenzie, K. J. et al. Ribonuclease H2 mutations induce a cGAS/STING-dependent innate immune response. EMBO J 35, 831–844, doi:10.15252/embj.201593339 (2016).

25 Behrendt, R. et al. Mouse SAMHD1 has antiretroviral activity and suppresses a spontaneous cell-intrinsic antiviral response. Cell Rep 4, 689–696, doi:10.1016/j.celrep.2013.07.037 (2013).

26 Ohto, T. et al. Intracellular virus sensor MDA5 mutation develops autoimmune myocarditis and nephritis. J Autoimmun 127, 102794, doi:10.1016/j.jaut.2022.102794 (2022).

27 Reijns, M. A. et al. Enzymatic removal of ribonucleotides from DNA is essential for mammalian genome integrity and development. Cell 149, 1008–1022, doi:10.1016/j.cell.2012.04.011 (2012).

28 Inoue, M. et al. An Aicardi-Goutieres Syndrome-Causative Point Mutation in Adar1 Gene Invokes Multiorgan Inflammation and Late-Onset Encephalopathy in Mice. J Immunol 207, 3016–3027, doi:10.4049/jimmunol.2100526 (2021).

29 Guo, X. et al. An AGS-associated mutation in ADAR1 catalytic domain results in early-onset and MDA5-dependent encephalopathy with IFN pathway activation in the brain. J Neuroinflammation 19, 285, doi:10.1186/s12974-022-02646-0 (2022).

30 Guo, X. et al. ADAR1 Zalpha domain P195A mutation activates the MDA5-dependent RNA-sensing signaling pathway in brain without decreasing overall RNA editing. Cell Rep 42, 112733, doi:10.1016/j.celrep.2023.112733 (2023).

31 Livingston, J. H. & Crow, Y. J. Neurologic Phenotypes Associated with Mutations in TREX1, RNASEH2A, RNASEH2B, RNASEH2C, SAMHD1, ADAR1, and IFIH1: Aicardi-Goutieres Syndrome and Beyond. Neuropediatrics 47, 355–360, doi:10.1055/s-0036-1592307 (2016).

32 Piccoli, C. et al. Late-Onset Aicardi-Goutieres Syndrome: A Characterization of Presenting Clinical Features. Pediatr Neurol 115, 1–6, doi:10.1016/j.pediatrneurol.2020.10.012 (2021).

33 D’Arrigo, S. et al. Aicardi-Goutieres syndrome: description of a late onset case. Dev Med Child Neurol 50, 631–634, doi:10.1111/j.1469-8749.2008.03033.x (2008).

34 Sun, L., Liu, S. & Chen, Z. J. SnapShot: pathways of antiviral innate immunity. Cell 140, 436–436 e432, doi:10.1016/j.cell.2010.01.041 (2010).

35 Kato, H. et al. Differential roles of MDA5 and RIG-I helicases in the recognition of RNA viruses. Nature 441, 101–105, doi:10.1038/nature04734 (2006).

36 Yoneyama, M. et al. Shared and unique functions of the DExD/H-box helicases RIG-I, MDA5, and LGP2 in antiviral innate immunity. J Immunol 175, 2851–2858 (2005).

37 Tang, Q. et al. Adenosine-to-inosine editing of endogenous Z-form RNA by the deaminase ADAR1 prevents spontaneous MAVS-dependent type I interferon responses. Immunity 54, 1961–1975 e1965, doi:10.1016/j.immuni.2021.08.011 (2021).

38 Nakahama, T. et al. Mutations in the adenosine deaminase ADAR1 that prevent endogenous Z-RNA binding induce Aicardi-Goutieres-syndrome-like encephalopathy. Immunity 54, 1976–1988 e1977, doi:10.1016/j.immuni.2021.08.022 (2021).

39 Maurano, M. et al. Protein kinase R and the integrated stress response drive immunopathology caused by mutations in the RNA deaminase ADAR1. Immunity 54, 1948–1960 e1945, doi:10.1016/j.immuni.2021.07.001 (2021).

40 de Reuver, R. et al. ADAR1 interaction with Z-RNA promotes editing of endogenous double-stranded RNA and prevents MDA5-dependent immune activation. Cell Rep 36, 109500, doi:10.1016/j.celrep.2021.109500 (2021).

41 Gitlin, L. et al. Essential role of mda-5 in type I IFN responses to polyriboinosinic:polyribocytidylic acid and encephalomyocarditis picornavirus. Proc Natl Acad Sci U S A 103, 8459–8464, doi:10.1073/pnas.0603082103 (2006).

42 Neeman, Y., Levanon, E. Y., Jantsch, M. F. & Eisenberg, E. RNA editing level in the mouse is determined by the genomic repeat repertoire. *RNA (New York*, N.Y 12, 1802–1809 (2006).

43 Levanon, E. Y. et al. Evolutionarily conserved human targets of adenosine to inosine RNA editing. Nucleic acids research 33, 1162–1168 (2005).

44 Eisenberg, E. et al. Is abundant A-to-I RNA editing primate-specific? Trends Genet 21, 77–81 (2005).

45. 45 Guo, X. L., S; Sheng, Y; Zenati, M; Billiar, T; Herbert A; Wang Q. ADAR1 Zα domain P195A mutation activates the MDA-5 dependent RNA-sensing signaling pathway in the brain without decreasing overall RNA editing. Cell Reports Accepted (2023).

46 Wang, Q. et al. Stress-induced apoptosis associated with null mutation of ADAR1 RNA editing deaminase gene. J Biol Chem 279, 4952–4961, doi:10.1074/jbc.M310162200 (2004).

